# Comparison of two sampling, quenching and extraction methods for quantitative yeasts metabolomics

**DOI:** 10.1101/2022.02.21.481278

**Authors:** Xiang Li, Mattia Rovetta, Patricia T.N. van Dam, Martin Pabst, Matthias Heinemann

## Abstract

Knowing intracellular metabolite concentrations is important for fundamental metabolism research as well as for biotechnology. The first steps in a quantitative metabolomics workflow, i.e., sampling, quenching, and extraction, are key to obtaining unbiased quantitative data. A qualified quenching and extraction method not only needs to rapidly terminate the *in vivo* biochemical reaction activities to preserve the endogenous metabolite levels but also has to fully extract all metabolites from cells. Recently, two different filtration-based sampling, quenching, and extraction methods have been proposed and used for quantitative yeast metabolomics. One method integrates the quenching and extraction into one step using a methanol-acetonitrile-water solution after a filtration step, while the other —more conventional— method quenches with cold methanol and extracts with boiling ethanol. In this study, we tested whether these two methods are equally well suited for quantitative metabolome analyses with yeast. Using isotope dilution mass spectrometry (IDMS) with GC-MS and LC-MS as analytical methods, in combination with precise quantification of cell volumes, we determined absolute concentrations of 63 intracellular metabolites covering amino acids, organic acids, phosphorylated sugars, and nucleotides in two *S. cerevisiae* strains with different physiology. By analyzing the data from samples generated with the two methods, we found that while both methods yielded essentially identical concentrations for amino and organic acids, the cold-solvent extraction yielded significantly lower concentrations for particularly phosphorylated sugars and nucleotides, presumably because of lower quenching or extraction efficiency of this method. Our results show that methanol-quenching combined with boiling-ethanol-extraction is the more accurate approach when aiming to quantify a broad range of different metabolites from yeast cells. The significant discrepancy observed between both common metabolite extraction methods demonstrates the importance of method optimization for quantitative studies in particular when working with microbes with rigid cell walls such as those found in yeast.

## INTRODUCTION

Metabolomics is one of the cornerstones in quantitative biology, which captures snapshots of intracellular metabolic status and provides insights into fundamental cellular processes [1,2]. For instance, metabolomics can facilitate unraveling the mechanisms behind cellular responses to changing environments or nutrient availability and can strengthen our understanding of the distribution of metabolic flux which provides vital information to identify bottlenecks in cell factories [3–5].

However, quantitative metabolomics is hindered by challenges such as the high turnover rates of metabolite pools, instability of certain metabolites during the sample preparation, and complex manipulation steps, which may introduce confounding variations and poor reducibility [6,7]. Quenching and extraction is the first step of the sample preparation in a quantitative metabolomics workflow and as such plays a crucial role in the quantification of intracellular metabolites levels. For accurate quantitative metabolite measurements, the quenching and extraction procedure is required to rapidly terminate all biochemical reaction activities while preserving the integrity of the metabolites and avoiding further degradation [5,8].

For yeast metabolomics, in the past, many different methods have been developed and been compared, which included the optimization of quenching solution and temperature, introducing washing steps, and altering the composition of the extraction reagents [1,3,7–9]. In recent years, two prevailing practices have surfaced. These are the cold-solvent (CS) quenching-extraction method and the cold methanol quenching-boiling ethanol extraction method (CMBE). Originally developed for metabolomics on microbes and further proven versatility for also mammalian cells and mouse tissues [3,10–12], the CS method exploits a fast filtration step and merges the quenching and extraction into one step by depositing the filter in a methanol-acetonitrile-water solution under a low temperature, aimed at minimizing the leakage losses of intracellular metabolites caused by the cold methanol quenching in both bacteria and yeast as reported [3,9,13]. In contrast, the CMBE method slices the workflow into three steps: first, quenching of cells using cold methanol, then collecting cells on a filter, and finally use of boiling ethanol to extract the metabolites [14]. The CMBE is proven efficient for extracting a wide scope of metabolites in yeast [8,9]. However, it turns out that a head-to-head comparison of the two methods has not been done and thus it is unclear whether they are equally suitable for quantitative metabolomics with yeast.

Here, to evaluate the performance of those two sampling, quenching, and extraction methods and to check their possible impacts on different metabolites, we applied the two methods on a *S. cerevisiae* wild-type strain and a hexose-transporters (*hxt*) mutant strain. Here, we found discrepancies in the total number of metabolites detected and in the concentrations of individual metabolites between these two methods.

## RESULTS

### Strains and cultivation

For our analyses, we used two yeast strains. One of them is a wild-type strain (WT), growing fermentatively, and the other one is a *hxt*-mutant strain (TM6*), growing with a respiratory metabolism. We grew our cultures in shake flasks, where we used an online gas transfer rates measurement device (TOM) to monitor the gas transfer rates (e.g., carbon dioxide transfer rate, CTR) in each shake flask [15]. This allowed us to (i) check for proper growth conditions and (ii) to assess cell growth in a minimally invasive manner. Specifically, we inoculated *S. cerevisiae* KOY wild-type strain and TM6* strain into different TOM flasks (biological triplicates for each strain). We collected the samples for our metabolomics analyses during the mid-exponential phase as reported by the carbon dioxide transfer rates (Supplementary Figure S1). Shortly before sampling for the metabolite analyses, we used a cell counter and analyzer CASY to determine both cell counts and cell size distributions. The average total cell volumes using the CS method were slightly lower than that of the CMBE method in both wild-type and TM6* strain (Supplementary Figure S2), as samples were always collected first to test the CS method then the CMBE method.

### Quenching and extraction workflow

To assess the two sampling-quenching-extraction methods, a parallel experiment was set up (Figure 1). In short, either the *S. cerevisiae* KOY WT strain or TM6* strain was inoculated in the glucose minimal medium and propagated to the mid-exponential phase and approximately 2~3 mg dry weight of cells was withdrawn from the culture. Then, the samples were processed with either of the two methods, until the concentration step. Using the CS method, the collected cell culture was immediately filtrated, and the filter was then transferred into the prechilled extraction solution (−20°C). After adding ^13^C-labeled yeast extract as an internal standard, metabolites were then extracted overnight at −20°C. Finally, the extraction solution was centrifuged to remove residual cell debris after the extraction, and the supernatant of samples was either stored at −70°C or dried to concentrate.

**Figure 1:**
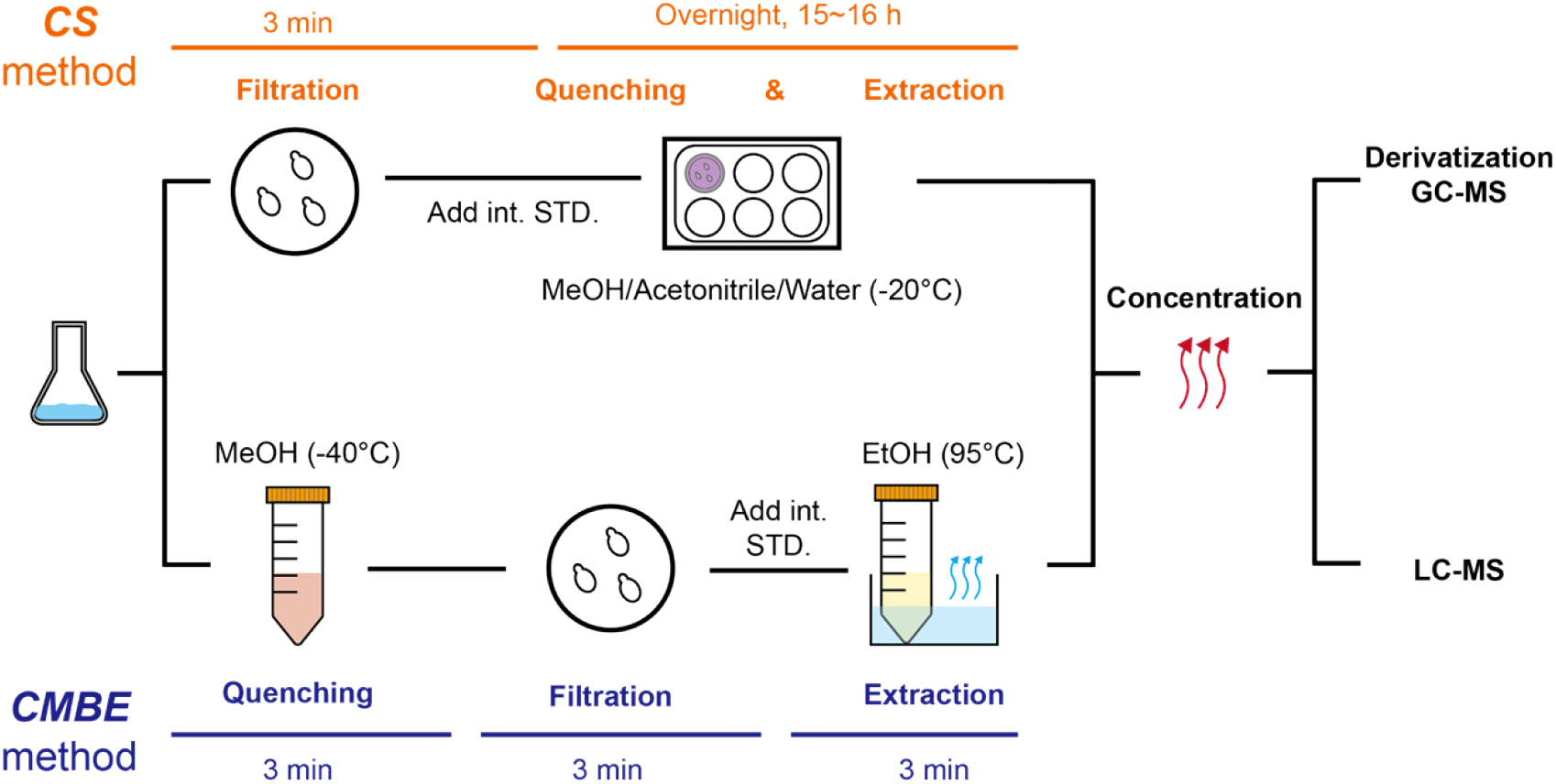
The workflow of the two sampling-quenching-extraction methods. CS method: <u>c</u>old-solvent quenching-extraction method; CMBE method: <u>c</u>old <u>m</u>ethanol quenching-<u>b</u>oiling <u>e</u>thanol extraction method; EtOH: ethanol; MeOH: methanol; int. STD: internal standard (^13^C-labeled yeast extract in this study); GC-MS: gas chromatography-mass spectrometry; LC-MS: liquid chromatography-mass spectrometry.

In the CMBE method, the metabolic activity was immediately quenched by adding cold methanol (−40°C) to the culture broth, and then this solution was filtrated to remove both medium and methanol. The filter containing the cell pellet was washed twice with pure methanol (−40°C) and immersed in a 75% (v/v) aqueous ethanol solution preheated to 75°C, to which again ^13^C-labeled internal standard was added. To extract the metabolites, cell pellets were boiled with ethanol at 95°C for three minutes.

The extract solutions from the two methods were dried using a RapidVap evaporator (Labconco) at 30°C for three hours and resuspended with 500 μL of MilliQ water. After removing the cell debris in the extract solution via centrifuging, the concentrations of amino acids, organic acids, and phosphorylated sugars were analyzed using GC-MS (except for fructose-1,6-biphosphate determined by LC-MS), whereas both nucleotides and coenzymes were determined by LC-MS.

### Comparison of metabolite concentrations obtained with both methods

We then compared the metabolite concentrations that we obtained from the two sampling, quenching, and extraction methods. To this end, we plotted the concentration of each metabolite extracted with the CS method against the one obtained from the CMBE method. For metabolites that lie on the diagonal, both methods yield the same results. Here, one can notice that with the exception of one metabolite (i.e., glucose) all metabolites lie either on the diagonal or under it (Figure 2A, 2B). Glucose is a special metabolite as it is not only present intracellularly, but it is also the growth substrate used in these experiments, and thus it is present in high concentrations in the medium. The fact that with both methods the determined glucose amount would account for unreasonably high intracellular concentrations (i.e. up to 78.59 ± 9.61 mM in WT and 168.20 ± 6.49 mM in TM6* with the CS method, which both concentrations even higher than the glucose concentration in the medium at the beginning of the batch culture (~ 55 mM)), this suggests that there is incomplete removal of the medium during the sample preparation or the carry-over effect as previously reported [16], which seems to be higher in the CS method (Figure 2A, 2B), likely due to the lack of a washing step in this method.

**Figure 2:**
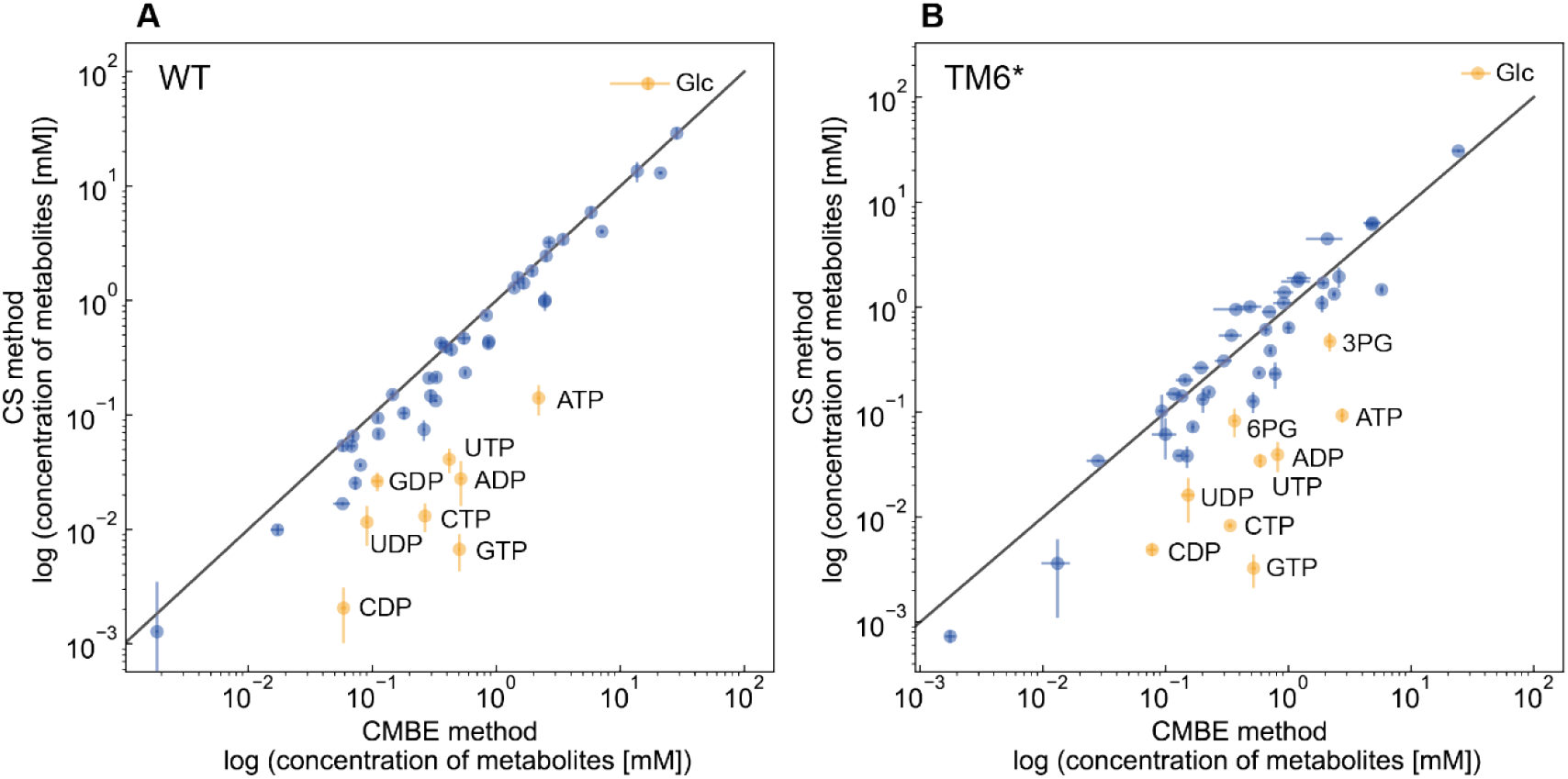
Comparison of intracellular concentrations of metabolites extracted with two methods. Mean concentrations of metabolites processed using the CS method were plotted against mean concentrations of corresponding metabolites derived from the CMBE method in a log-scale for both *S. cerevisiae* KOY WT strain (A) and KOY TM6* strain (B) with the respective standard deviations obtained from triplicates. The diagonal line indicates where metabolite concentrations are equal between both methods. For each data point, the distance from the point to the diagonal was calculated. Data points with a distance above 1 are labeled orange data points while data points close to (or on) the diagonal were labeled as blue. Outliers were annotated in the figures using their abbreviations, whose concentrations can be found in Supplementary Table S1.

Focusing on the remaining metabolites, we found that with the CMBE method we could quantify more metabolites. For instance, coenzymes such NAD^+^, NADP^+^, and CoA were undetectable using the CS method (Supplementary Table S1). Furthermore, it can be noticed that while a large number of metabolites essentially lie on the diagonal in the comparison plot, a number of metabolites also lie below (Figure 2A, 2B, Supplementary Table S1), meaning that the CS method yielded lower concentrations, indicating that metabolites either got lost or were insufficiently extracted with this method. These metabolites are mostly nucleotides and in the TM6* sample —in addition— also the glycolytic intermediates 3-phosphoglyceric acid (3PG) and 6-phosphogluconate (6PG) (Figure 2B). This suggests that the CS quenching-extraction method was either detrimental to both nucleotides and coenzymes or less efficient in extracting those metabolites.

### Quenching and extraction methods impacted metabolite classes differently

For a closer inspection of how those two methods impacted metabolites diversely, we first clustered intracellular metabolites into four classes based on their biochemical characteristics or associated pathways: amino acids, organic acids (TCA cycle), phosphorylated sugars (glycolysis, pentose phosphate pathway [PPP]), and nucleotides. For each metabolite and strain, we then performed a two-sided T-test among the triplicate samples quenched and extracted using the CMBE method and the other triplicates obtained from the CS method and plotted all *p*-values.

We found that the quenching and extraction methods impacted the classes of metabolites differently (Figure 3). For amino acids and organic acids (TCA) we obtained similar concentrations with both methods, as no significant differences were found in almost half of the candidates in the two groups (*p*-value > 0.05), while notably there are some marginal differences in the concentrations of some metabolites (e.g., Lys, Orn, and Cit). Consistent with the results of the globally lower nucleotides concentrations in the CS method compared to the CMBE method (Figure 2), nucleotides concentrations (except AMP in the TM6* strain) were severely different between the two methods (Figure 3). Also, phosphorylated sugars were sensitive to the selection of quenching and extraction methods as most *p*-values of candidates in this group were below 0.05.

**Figure 3:**
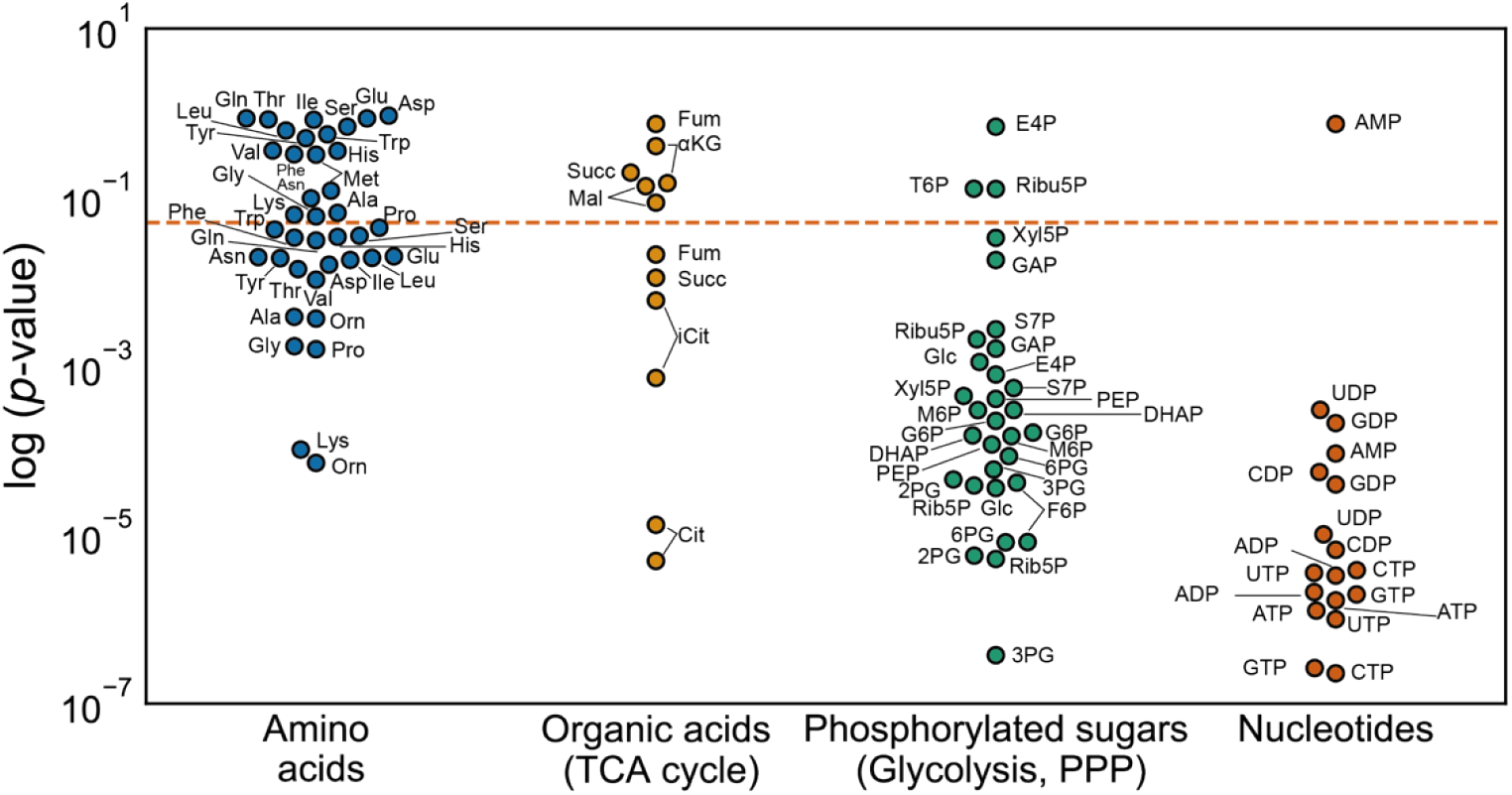
*p*-values of two-sided T-test for concentrations of metabolites extracted using the CS method and CMBE method. For each metabolite in each strain, a *p*-value was calculated using triplicate concentration measured by the CS method and another triplicate concentration from the CMBE method. *p*-values from both WT and TM6* strains were plotted together in log scale. Metabolites except coenzymes were clustered into four classes: amino acids, organic acids (TCA cycle), phosphorylated sugars (glycolysis, PPP), and nucleotides. The orange dashed line indicates a statistical significance threshold *p*-value of 0.05.

## DISCUSSION

Using two prevailing quenching-extraction methods in two yeast strains, we determined the absolute intracellular concentrations of metabolites. Interestingly, we found that different quenching and extraction methods can lead to metabolite class-dependent concentration biases. For instance, signals of coenzymes vanished, and levels of nucleotides and phosphorylated sugars were globally lower when using the CS method.

Consistent with previous reports, the cold-solvent (CS) method using a methanol-acetonitrile-water solution is likely to lead to systematically lower metabolite concentrations with phosphorylated sugars and nucleotides [8,9]. We propose three possible explanations for why the CS method underperformed compared to the more conventional CMBE method.

The first possible explanation for the observed lower concentrations of phosphorylated sugars and nucleotides in the CS method could be inefficient quenching. Compared to the CMBE method, which quenches metabolism using methanol at −40°C, the CS method will first collect cells via filtration at room temperature before the cell pellet is transferred into the quenching solution at −20°C. The duration of this filtration step depends on the sample volume, cell density, and viscosity of the culture, and here it took roughly three minutes for each sample. Considering the rapid turnover rates of some metabolites, proper temperature, and the timeframe during the filtration of unquenched culture could leave some space for intracellular reactions to proceed. If the quenching is not fully effective, we expect the adenylate energy charge (EC) to be lower as ATP is continued to be consumed. To test this, we calculated the energy charge in all samples (Supplementary Figure S3) and found that samples extracted using the CMBE method had consistent EC values of around 0.88 in both WT and TM6* strain, indicating that they had been quenched equally and sufficiently. However, processing the exact same cell samples with the CS method yielded generally lower EC values, with the EC values of TM6* even being as low as 0.5, hinting at insufficient quenching with the CS method.

Alternatively, it could also be envisioned that nucleotides are spontaneously interconverted into each other in the CS method. Emerging evidence suggests that the cellular matrix (defined as a set of total intracellular metabolites) could prevent the degradation of nucleotide triphosphates [17,18]. Though cellular matrix was present in both methods, considering the difference in the volumes of the extraction solution (4 mL in CS *vs.* 30 mL in CMBE), the cellular matrix extracted with the CMBE method was lower compared to the CS method. Therefore, we believe that this explanation of nucleotide interconversion is less likely as the cellular matrix was even more abundant in the CS method.

Another explanation could be the insufficient extraction in the CS method. In the CS method, the quenching and extraction are merged into one step, compared to the breaking up of cells using the boiling ethanol, the slower and milder extraction using the methanol-acetonitrile-water solution may not be enough to pull out all intracellular metabolites, even giving a long extraction period (15~16 hours) at low temperature (−20°C). Previously, it was suggested that even though cell lysis is complete using the CS method [8], polar (or charged) metabolites could be partially insoluble in this cold organic solvent, and they could be missed out partly if they are removed through precipitation with cell debris during the later washing steps [8]. We consider this reasonable as we found snowflake-like aggregates of cell debris in the multi-well plate after the overnight extraction, and we can imagine that not completely soluble phosphorylated sugars and nucleotides could be washed down with the cell debris.

Taken together, our findings demonstrate that particularly concentrations of coenzymes, phosphorylated sugars, and nucleotides are subject to the choice of sampling-quenching-extraction methods. Our verdict is that the conventional methanol-quenching and boiling-ethanol-extraction (CMBE) method should be preferably used compared to the cold-solvent quenching-extraction (CS) method for quantitative yeast metabolomics studies. One should be cautious when choosing sample preparation steps for yeast metabolomics as systematic errors may bias overall concentrations or introduce a bias to specific classes of metabolites.

## MATERIAL AND METHODS

### Yeast strains and maintenance

*S. cerevisiae* KOY PK2-1C83 (wild type) strain and its mutant strain TM6* were used in this study [19]. Both yeast strains were propagated in YPD medium (1% [w/v] yeast extract, 2% [w/v] peptone, 2 % [w/v] glucose) at 30°C. Before the main experiment, both glycerol stocks were streaked on the YPD agar plate and incubated at 30°C to obtain single colonies.

### Shake flask cultivation

All batch cultures in this study were performed in the minimal medium [20] supplemented with 10 g·L^−1^ glucose at 30°C and a shaking speed of 300 rpm. To ensure cells were in optimal physiological conditions, a two-step pre-culture strategy was followed. Single colonies of each strain were transferred from the YPD agar plate and inoculated in 10 mL of minimal medium (pH 5.0, 10 g·L^−1^ glucose) in the first batch of the pre-culture for eight hours. The first batch of the pre-culture was subsequently diluted into another 10 mL of preheated minimal medium (30°C) for the second step of pre-culture to reach a cell density OD_600_ of 1~2 (mid-exponential phase) after the overnight cultivation. Then, the overnight pre-culture was diluted to preheated 50 mL minimal medium (10 g·L^−1^ glucose) with a starting cell count between 1×10^6^ and 2×10^6^ in 500 mL flasks, which were connected to an online gas transfer rates monitoring device TOM (Kuhner GmbH). The growth of cells was followed by monitoring their carbon dioxide transfer rates (CTR) and oxygen transfer rates (OTR), which, in an exponentially growing culture, both scales with cell counts in the culture. For each strain, three replicates were inoculated and sampled.

### Quantification of the total cell volume

To determine the total cell volume of each sample, a cell counter and analyzer CASY (OLS OMNI Life Science) was applied to measure both cell counts and cell size distributions. Right before quenching and extraction, an aliquot of 50 μL of cell culture was withdrawn from the shaking flask and thoroughly mixed with 10 mL of CASY TON solution before loading on the cell counter CASY. For each sample, the CASY measurements return a value of a number of cells (*n_d_*) in each diameter (*d*) channel, from which the total cell volume (*V_tot_*) was calculated by integrating cell volumes in each diameter channel from 0 to 20 μm, and then amplified back by multiplying a dilution constant *D*:

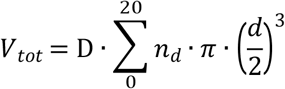

### CMBE method: rapid sampling with cold methanol quenching and boiling ethanol extraction

Before sampling, the optical density of the exponentially growing cultures was determined via a spectrophotometer. By the moment when OD_600_ of the culture reached 1.5 ~ 2.0, approximately 5 to 6 mL broth (equivalent to 2~3 mg of dry biomass) was withdrawn from each culture and immediately quenched by adding to an aliquot of 25 mL of cold methanol (−40°C). The quenched culture-methanol mixture was immediately vortexed for 10 seconds and filtrated through a biomass filter (Supor-200, 0.2 μm, 47 mm, Pall Corporation), which was pre-layered with 15 mL of pure methanol (−40°C). The filter containing the quenched and filtered biomass was subsequently washed with another 15 mL of pure methanol (−40°C) and immediately transferred into a 50-mL Falcon tube containing 30 mL of preheated (75°C) ethanol solution (75% v/v). After spiking an aliquot of 100 μL of ^13^C-labeled cell extract into the ethanol solution as an internal standard [21,22], quenched cells were extracted at 95°C for 3 minutes using a water bath and chilled on ice afterward. When the extraction was done, the biomass filter was removed, and the extraction solution was stored at −70°C before later processing. The remaining ^13^C-labeled cell extract was also preserved at −70°C for creating calibration curves during metabolite analysis. To further concentrate analytes, extraction solutions were dehydrated at 30°C using a RapidVap Dry Evaporator (Labconco) following the standard procedure for three hours. The dried sediment in extracted samples was then resuspended in an aliquot of 500 μL MilliQ water, vortexed, and transferred into (numbered) 1.5-mL Eppendorf tubes. After centrifuging at 13,000 rpm for 5 minutes at 1°C, the supernatant was collected and centrifuged again using the same centrifuge settings. The supernatant was eventually stored at −80°C until further analysis.

### CS method: integrated quenching and extraction with cold methanol-acetonitrile-water solution

A day prior to sampling, the extraction solution (40:20:20 [%, v/v] methanol: acetonitrile: water, all in MS-grade) was prepared and stored in a six-well plate (4 mL per well) at −20°C. Before sampling, the OD_600_ of the culture was monitored till the OD_600_ reached 1.5~2, and then a certain amount of culture (2~3 mg dry weight) was filtrated through a biomass filter (Durapore 0.45 μm, diameter 25-mm, Sigma). The filter with the cell pellet was immediately transferred to an aliquot of 4 mL of precooled extraction solution (−20°C) with an addition of an aliquot of 100 μL of ^13^C-labeled yeast extract and incubated at −20°C for overnight (15~16 hours). After the overnight extraction, the extract (~ 4 mL) was transferred into a 15-mL Falcon tube (prechilled on ice) and centrifuged for 5 minutes at −9°C at the maximum speed to remove the remaining cell debris. An aliquot of 3.5 mL supernatant was then transferred into another pre-labeled 15-mL Falcon tube and stored at −70°C. All samples were further concentrated as described in the CMBE method before being analyzed by LC-MS or GC-MS.

### Analysis of nucleotides by liquid chromatography-mass spectrometry

Identification and quantification of nucleotides were performed using an ACQUITY UPLC chromatography system (Waters, UK) coupled online to a high-resolution Orbitrap mass spectrometer (Q-Exactive Focus, Thermo Fisher Scientific, Germany). For chromatographic separation, a reverse phase (Kinetex C18 column 2.1 x 150 mm, 1.7 μm; Phenomenex, Torrance, CA, USA)[23] was used at room temperature using 50 mM ammonium acetate pH 5.0 as mobile phase A, and 90 % acetonitrile plus 10% 50 mM ammonium acetate pH 5.0 acid as mobile phase B (v/v). A flow of 250 μL·min^−1^ was maintained over a solvent composition from 50% to 75% B over 4 minutes, before equilibrating back to the starting conditions. The metabolite extracts (spiked with the ^13^C yeast metabolite extract) were taken from −80°C and brought to room temperature immediately before injection. The samples were carefully vortexed and 5 μL were subsequently injected onto the UPLC reverse phase separation system [24]. The mass spectrometer was operated in negative ionization mode, where a full scan from 250‒1000 m/z was acquired in ESI positive mode (−2.8 kV) at a resolution of 70 K, an AGC target of 1e6, and by setting the max injection time to auto. The ^13^C metabolite yeast extract and synthetic standards from each metabolite were analyzed additionally in separate runs to confirm elution order, accurate mass, and main fragments. Mass spectrometric raw data were analyzed using XCalibur 4.1 (Thermo) and after conversion to mzXML files by MsConvert using Matlab 2020b, where the peak intensities of the individual metabolite compounds were summed and expressed as intensity ratios to their corresponding ^13^C yeast extract metabolite peaks (internal standard). Absolute quantification was based on an external calibration curve established from commercial standards spiked with the same ^13^C labeled yeast metabolite extract [21,24,25]. The mass spectrometer was calibrated using the Pierce™ LTQ negative ion calibration solution (Thermo Fisher Scientific, Germany).

### Analysis of sugars and amino acids by gas chromatography-mass spectrometry

For the analysis of sugars and sugar phosphates an aliquot of 100 μL concentrated extract was lyophilized and resuspended in 50 μL of 20 mg·mL^−1^ methoxylamine HCl (in pyridine) solution. After the incubation of 50 minutes at 70°C, an aliquot of 80 μL of MSTFA/TMCS (20:1 v/v, Sigma) was added to each sample, and samples were incubated at 70°C for another 50 minutes.

For the analysis of amino acids 30 μL of 10 mg·mL^−1^ NaCl was added to an aliquot of 100 μL concentrated extract. The solution was lyophilized, then resuspended in 75 μL of acetonitrile and 75 μL of MTBSTFA and incubated at 70°C for 60 minutes. Identification and quantification of the compounds were essentially performed as described [22,26].

### Statistical analysis

Using a python package Scipy (Python version 3.7.4, Scipy version 1.7.3), a two-sided student T-test was performed to evaluate the impacts of different quenching-extraction methods on different species of metabolites. To determine whether concentrations measured from one method were significantly different from the other, for each metabolite in each strain, a *p*-value was calculated using triplicate concentration measured by the CS method and another triplicate concentration from the CMBE method. The results of the T-test could be found in Supplementary Table S2.

## ACKNOWLEDGEMENT

The work of XL is funded by China Scholarship Council (No. 201806790008).

## AUTHOR CONTRIBUTION

XL, MR, and MH conceived and designed the research; XL, MR, and PD performed the experiments; XL, PD, MP, and MH analyzed data; MH supervised the project. XL and MH drafted the manuscript. All authors read and approved the final manuscript.

## Supplementary files

**Supplementary Figure S1:**
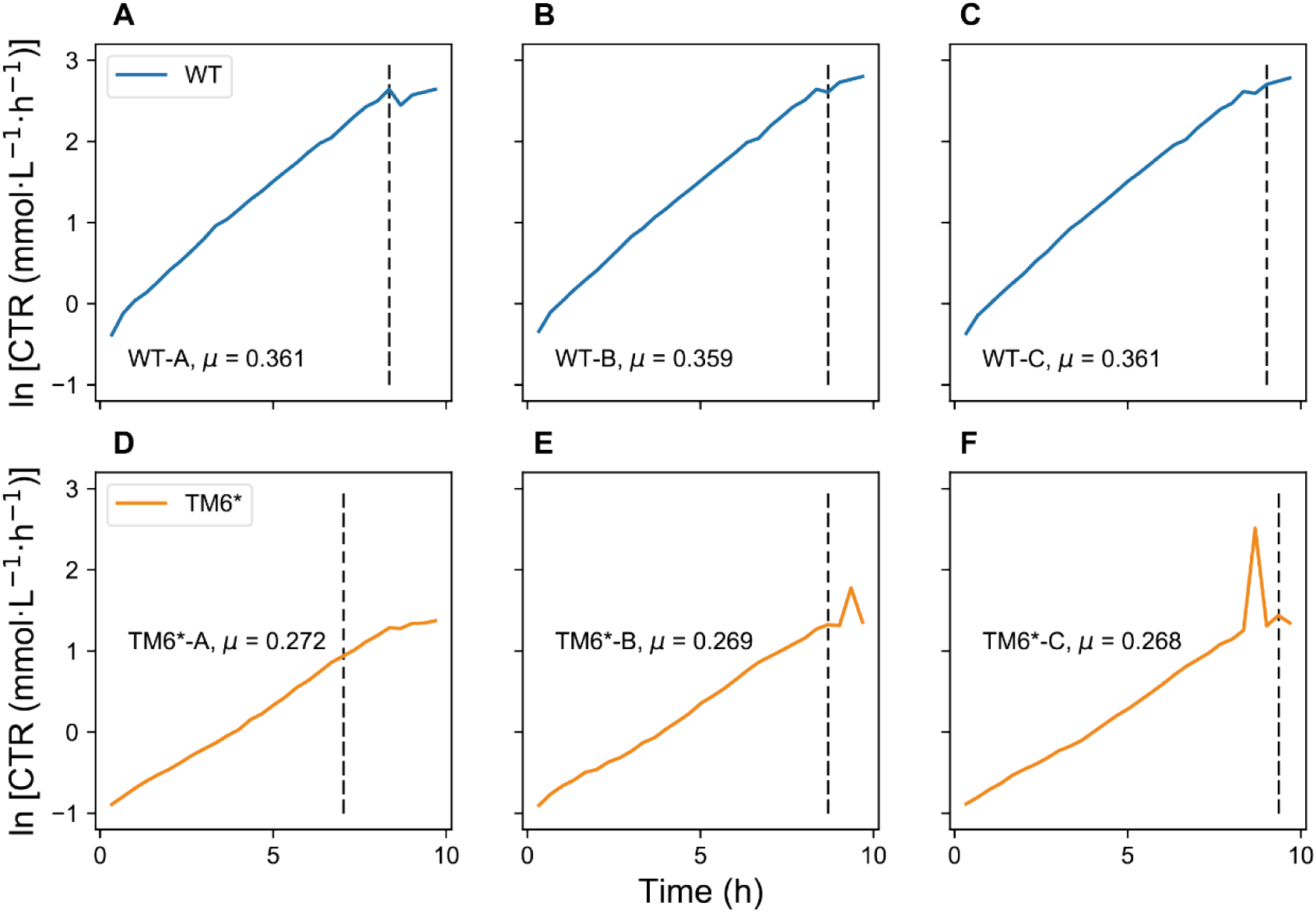
Growth of each sample monitored by its carbon dioxide transfer rate (CTR) in each flask. Figure A-C represents the triplicate of *S. cerevisiae* KOY wild-type strain and figures D-F shows the cell propagation of triplicate of *S. cerevisiae* KOY TM6* strain. After transforming the CTR of all samples into the log scale, a linear regression was performed for each sample, and their approximate growth rates before sampling were labeled on the figure.

**Supplementary Figure S2:**
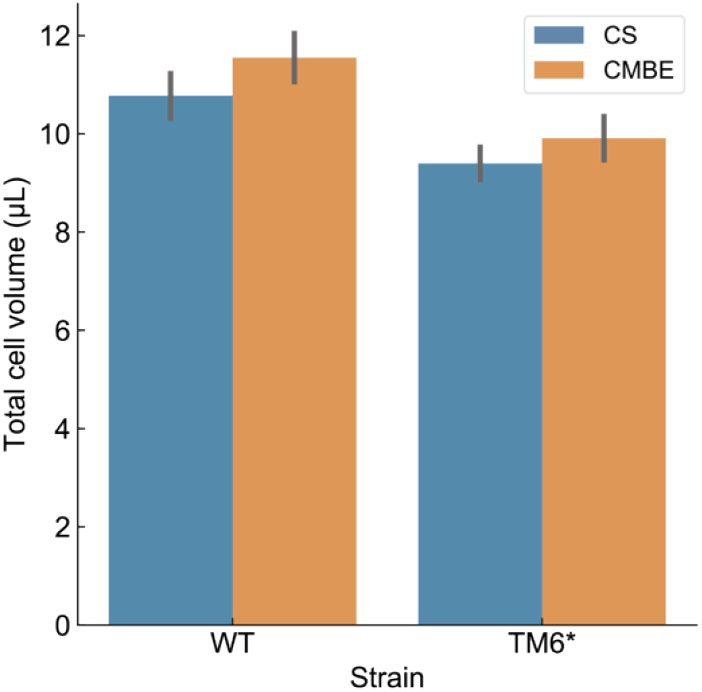
Average total cell volume in samples among different strains and methods. For each strain, biological triplicate samples were prepared to test the CS method and CMBE method individually, and the total cell volume (μL) in each sample was measured and calculated using the cell counter and analyzer CASY device. Blue bars indicate the average total cell volume of samples used to test the CS method, while orange bars indicate the average total cell volume of samples used to test the CMBE method.

**Supplementary Figure S3:**
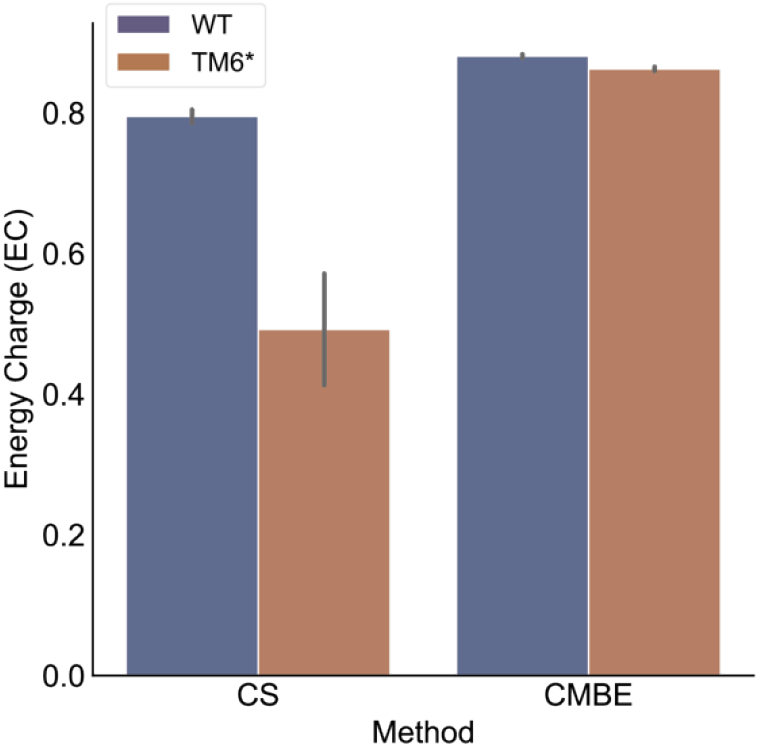
Adenylate energy charge (EC) in different groups of samples. For each strain, biological triplicate samples were prepared to test the CS method and CMBE method individually, and the adenylate energy charge (EC) was calculated using intracellular concentrations of AMP, ADP, and ATP in each sample. Dark blue bars indicate the average EC calculated in the WT strain, while the dark orange bars indicate the average EC calculated from the TM6* strain.

